# Significance of linkage disequilibrium and epistasis on the genetic variances and covariance between relatives in non-inbred and inbred populations

**DOI:** 10.1101/2021.01.19.427275

**Authors:** José Marcelo Soriano Viana, Antonio Augusto Franco Garcia

## Abstract

Because no feasible theoretical model can depict the complexity of phenotype development from a genotype, the joint significance of linkage disequilibrium (LD), epistasis, and inbreeding on the genetic variances remains unclear. The objective of this investigation was to assess the impact of LD and epistasis on the genetic variances and covariances between relatives in non-inbred and inbred populations using simulated data. We provided the theoretical background and simulated grain yield assuming 400 genes in 10 chromosomes of 200 and 50 cM. We generated five populations with low to high LD levels, assuming 10 generations of random cross and selfing. The analysis of the parametric LD in the populations shows that the LD level depends mainly on the gene density. The significance of the LD level is impressive on the magnitude of the genotypic and additive variances, which is the most important component of the genotypic variance, regardless of the LD level and the degree of inbreeding. Regardless of the type of epistasis, the ratio epistatic variance/genotypic variance is proportional to the percentage of the epistatic genes. For the epistatic variances, except for duplicate epistasis and dominant and recessive epistasis, with 100% of epistatic genes, their magnitudes are much lower than the magnitude of the additive variance. The additive x additive variance is the most important epistatic variance. Our results explain why LD for genes and relationship information are key factors affecting the genomic prediction accuracy of complex traits and the efficacy of association studies.

## Introduction

Genomic selection or genomic prediction of complex quantitative traits and genome-wide association studies (GWAS) share a common quantitative genetics background: linkage disequilibrium (LD) between genes and single nucleotide polymorphisms (SNPs). Among the thousands of studies that have been published in these areas, only a few have offered a quantitative genetics background, indicating that the accuracy of the genomic prediction and the power to detect a candidate gene depends on the LD between genes and SNPs (Gianola et al. 2009; Goddard 2009; Viana et al. 2016, 2017a). Based on Cockerham (1954), Viana et al. (2016) and Viana et al. (2017a) provided explicit functions relating SNP additive, dominance, and epistatic effects and variances to the quantitative trait locus (QTL) additive, dominance, and epistatic effects and variances. For a biallelic QTL (alleles A/a) and a SNP (alleles B/b) in LD, the connection between both their effects and variances is the measure of LD in the gametic pool of the population, which is given by the difference between the products of the haplotypes Δ = *P_AB_ P_ab_ - P_Ab_P_aB_* (Kempthorne, 1973). This LD measure can be expressed as 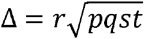 and Δ = *D*’.max (*e*), where r is the correlation between the values of alleles in both loci in the same gamete, p and q are the QTL allelic frequencies, s and t are the SNP allelic frequencies, D’ is a relative measure of LD (in relation to the maximum value for given allele frequencies), and max(e) is the maximum deviation of the actual gamete frequency from linkage equilibrium (Hill and Robertson 1968; Lewontin 1964).

However, the relationship information is the most important factor affecting the genomic prediction accuracy. For GWAS, the relationship information allows an effective control of the type I error rate and decreases the number of significant associations outside of the QTL intervals (Liu et al. 2015; Pereira et al. 2018). Although the observed relationship information, which is computed from the molecular data, has provided superior prediction accuracies and a more effective control of the false discovery rate (FDR), relative to the expected relationship information (Liu et al. 2016; Munoz et al. 2014), the covariance between relatives is another significant quantitative genetics background in the quantitative genomics era.

Several investigations on genomic selection, genomic prediction of complex traits, and GWAS have included epistasis (Monir and Zhu 2017; Varona et al. 2018). With few exceptions, a common feature of these significant studies is the absence of basic genetics background on the epistasis types (see any standard genetics book). These studies consider the basic quantitative genetics background for modelling epistasis, but there is generally no reference for the key knowledge provided by Kempthorne (1954) and Cockerham (1954) on modelling epistatic effects and defining the epistatic genomic relationship matrices. That is, for Bayesian and random regression (based on SNP effects) and genomic (based on genetic effects) best linear unbiased prediction (BLUP) approaches, the additive x additive, additive x dominance, dominance x additive, and dominance x dominance SNP/QTL effects are modeled, but in simulation-based studies there is no specification of any type of epistasis as complementary, duplicate, dominant, recessive, dominant and recessive, duplicate genes with cumulative effects, and non-epistatic genic interaction. The significant scientific contributions of these studies rely on the definition of SNP coding for the epistatic effects and specification of the epistatic genomic relationship matrices, which are deduced following the key proposition by VanRaden (2008), aiming maximization of the prediction accuracy and power to detect interacting QTLs (Jiang and Reif 2020; Martini et al. 2017; Vitezica et al. 2017).

Because of negative consequences (nominated inbreeding depression), human, conservative, animal, and cross-pollinated species geneticists agree that inbreeding should be efficiently controlled to maintain adequate genetic diversity in the populations (Hasselgren and Noren 2019; Howard et al. 2017). However, self-pollination has been deliberately used in maize hybrid breeding (currently to a lesser extent due to the doubled-haploid technology). For self-pollinated crops, the development of varieties involves selection over generations of increasing inbreeding. In these populations the inbreeding has an impact on the genetic variances and covariance between relatives (Cockerham 1983).

As previously highlighted, the most important quantitative genetics theory for modelling epistasis was developed by Kempthorne (1954). Cockerham (1954) also provided a significant contribution. If modelling only inbreeding, LD, or epistasis is a difficult task for the quantitative geneticists, jointly modelling the three events is a challenge. An impressive approach for two genes theory in quantitative genetics assuming inbreeding, LD, and epistasis was presented by Weir and Cockerham (1977). Because of the complexity of the expressions for the genetic variances and covariance between relatives, they concluded that “the result is of little use”. That is, the functions do not allow assessing the influence of these factors on the genetic variability and the degree of relationship in the populations. Because LD, relationship information, epistasis, and inbreeding are key factors for genomic prediction and association studies, the objective of this investigation was to assess the impact of LD and epistasis on the genetic variances and covariance between relatives in non-inbred and inbred populations, using a simulated data set.

## Material and Methods

### Additive and dominance genetic values in inbred populations in LD

Assume initially a single biallelic gene (A/a) determining a quantitative trait, where A is the gene that increases the trait expression. The genotype probabilities in a population derived by n generations of selfing from a Hardy-Weinberg equilibrium population (generation 0) are *p*^2^ + *pqF*, 2*pq*(1 − *F*), and *q*^2^ + *pqF*, if AA, Aa, and aa, respectively, where p and q are the allelic frequencies for A and a, respectively, and *F* = 1 − (1/2)^*n*^ is the inbreeding coefficient. The inbred population mean is *M_F_* = *m* + (*p − q*)*a* + 2*pqd* − 2*Fpqd* = *M* − 2*Fpqd*, where m is the mean of the genotypic values of the homozygotes, a is the difference between the genotypic value of the homozygote of greater expression and m, d is the dominance deviation, – *2Fpqd* is the change in the population mean due to inbreeding, and M is the mean of the non-inbred population (Hallauer and Miranda Filho, 1988). Defining 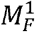 and 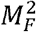 as the means of the inbred population after an allelic substitution for the genes A and a, respectively, the average effect of the allelic genes in the inbred population are 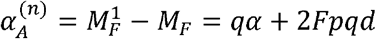 and 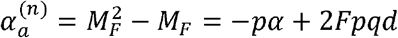 where 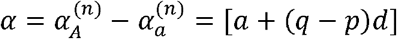 is the average effect of an allelic substitution. Note that the average effect of an allelic substitution is the same in the non-inbred and in the inbred populations. Thus, the additive values in the inbred population are 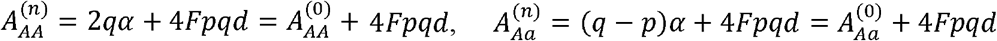 and 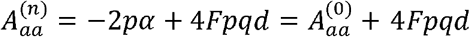 where *A*^(0)^ is the additive value in the non-inbred population. Note that *E*(*A*^(n)^) = 4*Fpqd*. Expressing the genotypic values in the inbred population as a function of *M_F_*, we have:

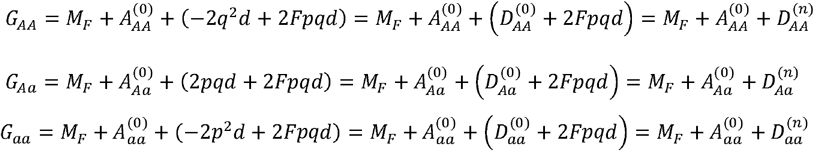

Note that in the inbred population, *E*(*A*^(0)^) = *E*(*D*^(n)^) = 0 but *E*(*D*^(0)^) = −2*Fpqd*. Note also that the additive value in the non-inbred population is the additive value in the inbred population expressed as deviation from its mean (*A*^(0)^ = *A*^(n)^ − 4*Fpqd*) and the dominance value in the inbred population is the dominance value in the non-inbred population expressed as deviation from its mean (*D*^(n)^ = *D*^(0)^ + 2*Fpqd*). This imply that, in the inbred population, *E*(*G*) = *M_F_*.

### Genetic variances in inbred populations in LD

Assume now two linked biallelic genes (A/a and B/b) determining a quantitative trait and a non-inbred population in LD (generation 0). Assume dominance but initially no epistasis. The genotype probabilities in generation 0 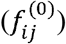 are presented by Viana (2004), where i and j (i, j = 0, 1, or 2) are the number of copies of the gene that increase the trait expression (A and B). For example. 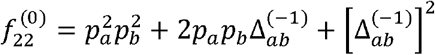, where 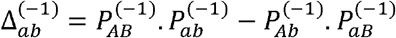 is the measure of LD in the gametic pool of generation −1 (Kempthorne, 1973). After n generations of selfing, the genotype probabilities are:

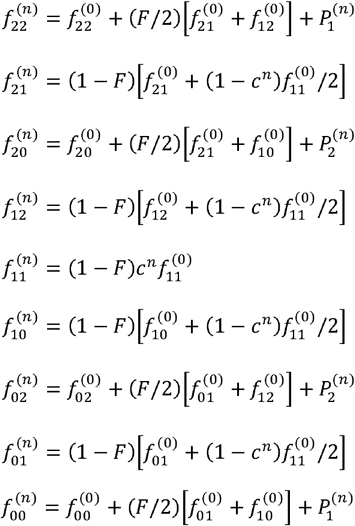

where

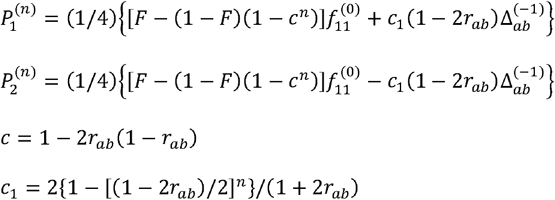

and *r_ab_* is the recombination frequency.

The genotypic variance for the two genes in the inbred population is 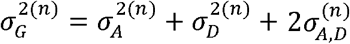, where:

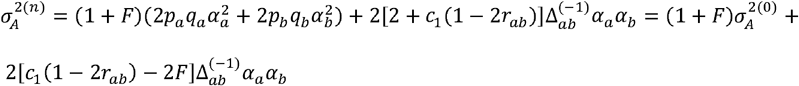

is the additive variance,

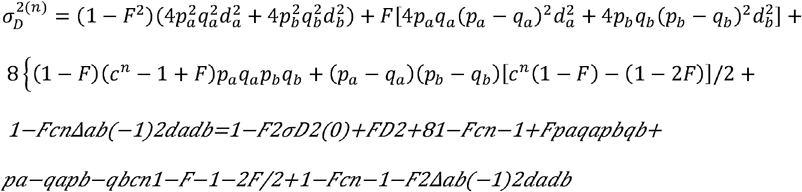

is the dominance variance, and

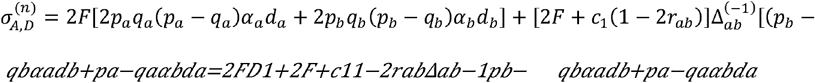

is the covariance between additive and dominance values,
where 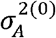 and 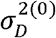 are the additive and dominance variances in the non-inbred population in LD (Viana 2004), and *D*_1_ (covariance of a and d) and *D*_2_ (variance of d) are the components of the covariance of relatives from self-fertilization, assuming linkage equilibrium (Cockerham 1983). The other terms are the covariances between the average effects of an allelic substitution, between dominance deviations, and between the average effect of an allelic substitution and dominance deviation, for genes in LD. Because we assumed biallelic genes, 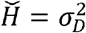. Thus, 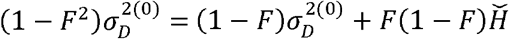. Note that the genotypic variance derived here is a general formulation for the Cockerham’s genotypic variance c_ggg_, assuming LD. If p = q, 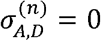.

Assuming LD but no inbreeding, the genotypic variance after n generations of random cross in the non-inbred population in LD is 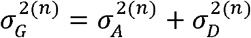, because 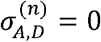, where:

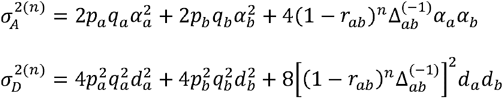

Thus, the genotypic variance decreases after n generations of random cross in a non-inbred population. Assuming LD, no inbreeding, and no epistasis, a general formulation for the covariance between relatives is

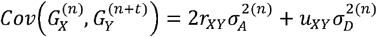

where *r_XY_* is the Malecot’s coefficient of relationship between the relatives X in generation n and Y in generation n + t and *u_XY_* is the probability that the parents of X (f and m, for the female and male) and Y (f’ and m’, for the female and male) have genotypes identical by descent (*u_X,Y_ = r_ff′_. r_mm′_ + r_fm′_ r_mf′_*). The covariance between parent and offspring and between half- and full-sibs were derived by Viana (2004):

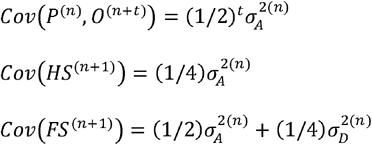

Assuming LD, inbreeding, and no epistasis, it is not straightforward to develop a general expression for the covariance between relatives from selfing (c_tgg’_) derived by Cockerham (1983), assuming no LD and no epistasis. Note that the covariance between parent (generation 0) and its selfed progeny (generation 1) derived by Viana (2004) depends also on the covariance between α and d for distinct genes in LD, but the coefficients for the additive and dominance variances and for *D*_1_ are those predicted by Cockerham (1983):

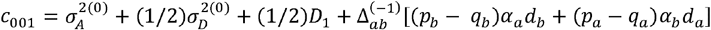

### Epistasis in non-inbred and inbred populations in LD

The quantitative genetics theory for modelling epistasis in a population in LD is a generalization of the theory proposed by Kempthorne (1954), who assumed a non-inbred population in linkage equilibrium and any number of alleles. We assumed biallelism. It should be highlighted that the Kempthorne’s theory allows a generalization from two to three or more interacting genes. But fitting three or more interacting genes in a population in LD is a challenge because the genotype probabilities for three or more genes in LD are too complex to derive. Furthermore, only complementary and duplicate epistasis can be easily defined for three or more epistatic genes.

Assume now that the two previous defined genes are epistatic. The genotypic value is (Kempthorne 1954):

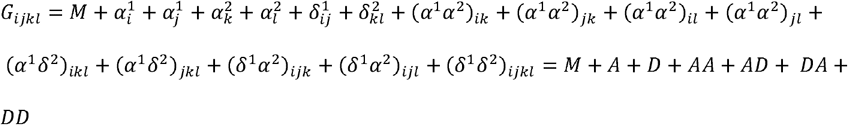

where AA, AD, DA, and DD are the additive x additive, additive x dominance, dominance x additive, and dominance x dominance epistatic genetic values.

The parametric values of the 36 parameters for the nine genotypic values are obtained by solving the equations *β* = (*X’VX)^-1^ X’Vy*, where *X* is the incidence matrix, 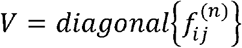 is the diagonal matrix of the genotype probabilities, and *y* is the vector of the genotypic values (*G_ij_*) (*i, j* = 0, 1, and 2). Following Kempthorne (1954), we completed the rank of X with 31 of the following restrictions (not all linearly independent):

i) restrictions for the average effects of genes: 1) *p_a_α_A_ + q_a_α_a_* = 0 and 2) *p_b_α_B_ + q_b_α_b_*.
ii) restrictions for the dominance values: 1) *p_a_δ_AA_ + q_a_δ_Aa_* = 0; 2) *p_a_δ_Aa_ + q_a_δ_aa_* = 0; 3) *P_b_δ_BB_* + *q_b_δ_Bb_* = 0, and 4) *P_b_δ_Bb_* + *q_b_δ_bb_* = 0.
iii) restrictions for the AA values: 1) *p_b_*(*α_A_α_B_*) + *q_b_*(*α_A_α_b_*) = 0; 2) *p_b_*(*α_a_α_B_*) + *q_b_*(*α_a_α_b_*) = 0; 3) *p_a_*(*α_A_α_B_*) + *q_a_*(*α_a_α_B_*) = 0; and 4) *p_a_*(*α_A_α_b_*) + *q_a_*(*α_a_α_b_*) = 0.
iv) restrictions for the AD values: 1) *p_b_*(*α_A_ δ_BB_*) + *q_b_*(*α_A_ δ_Bb_*) = 0; 2) *p_b_*(*α_A_δ_Bb_*) + *q_b_*(*α_A_δ_bb_*) = 0; 3) *p_b_*(*α_a_δ_BB_*) + *q_b_*(*α_a_δ_Bb_*) 0; 4) *P_b_*(*α_a_δ_Bb_*) + *q_b_*(*α_a_δ_bb_*) = 0; 5) *P_a_*(*α_A_δ_BB_*) + *q_a_*(*α_a_δ_BB_*) = 0; 6) *p_a_*(*α_A_δ_Bb_*) + *q_a_*(*α_a_δ_Bb_*) = 0; and 7) *p_a_*(*α_A_δ_bb_*) + *q_a_*(*α_a_δ_bb_*) = 0 (six out of the seven; four out of the seven if there is no dominance for the second gene).
v) restrictions for the DA values: 1) *p_a_*(*δ_AA_α_B_*) + *q_a_*(*δ_Aa_α_B_*) = 0; 2) *p_a_*(*δ_Aa_α_B_*) + *q_a_*(*δ_aa_α_B_*) = 3) *p_a_*(*δ_AA_α_b_*) + *q_a_*(*δ_Aa_α_b_*) = 0; 4) *p_a_*(*δ_AA_α_b_*) + *q_a_*(*δ_aa_α_b_*) = 0; 5) *p_b_*(*δ_AA_α_B_*) + *q_b_*(*δ_AA_α_b_*) = 0; 6) *p_b_*(*δ_Aa_α_B_*) + *q_b_*(*δ_Aa_α_b_*) = 0; 7) *p_b_*(*δ_aa_α_B_*) + *q_b_*(*δ_aa_α_b_*) = 0; (six out of the seven; four out of the seven if there is no dominance for the first gene).
vi) restrictions for the DD values: 1) *p_b_*(*δ_AA_δ_BB_*) + *q_b_*(*δ_AA_δ_Bb_*) = 0; 2) *p_b_*(*δ_AA_δ_Bb_*) + *q_b_*(*δ_AA_δ_bb_*) = 0; 3) *p_b_*(*δ_Aa_δ_BB_*) + *q_b_*(*δ_Aa_δ_Bb_*) = 0; 4) *p_b_*(*δ_Aa_δ_Bb_*) + *q_b_*(*δ_Aa_δ_bb_*) 0; 5) *P_b_*(*δ_aa_δ_BB_*) + *q_b_*(*δ_aa_δ_Bb_*) = 0; 6) *P_b_*(*δ_aa_δ_Bb_*) + *q_b_*(*δ_aa_δ_bb_*) = 0; 7) *P_a_*(*δ_AA_δ_BB_*) + *q_a_*(*δ_Aa_δ_BB_*) = 0; 8) *P_a_*(*δ_Aa_δ_BB_*) ; *q_a_*(*δ_aa_δ_BB_*) = 0; 9) *P_a_*(*δ_AA_δ_Bb_*) + *q_a_*(*δ_Aa_δ_Bb_*) = 0; 10) *p_a_*(*δ_Aa_δ_Bb_*) + *q_a_*(*δ_aa_δ_Bb_*) = 0; 11) *p_a_*(*δ_AA_δ_bb_*) + *q_a_*(*δ_Aa_δ_bb_*) = 0; and 12) *p_a_*(*δ_Aa_δ_bb_*) + *q_a_*(*δ_aa_δ_bb_*) = 0 (nine out of the 12; eight out of the 12 if there is no dominance for both genes).

Kempthorne (1954) provided explicit functions for all effects because he assumed linkage equilibrium. Assuming LD makes very difficult to derive such functions but the following results hold:

1. the expectation of the breeding value is zero regardless of the degree of inbreeding in the population.
2. the expectation of the dominance value is *E*(*D*)^(n)^ = *p_a_q_a_F*(*δ_AA_* - 2*δ_Aa_*+*δ_aa_*) + *p_b_q_b_F*(*δ_BB_* — 2*δ_Bb_*+*δ_bb_*); then, defining the dominance value in an inbred population as the dominance value expressed as deviation from its mean (*D^(n)^* = *D* — *E*(*D^(n)^*), *E*(*D^(n)^*) = 0.
3. the expectation of the additive x additive value is

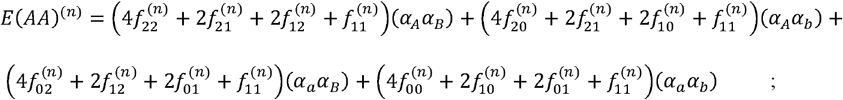

this expectation is zero only if there is no LD.
4. the expectation of the additive x dominance value is

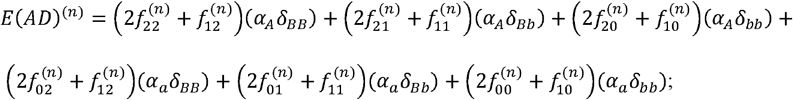

this expectation is zero only if F = 0 or if there is no LD.
5. the expectation of the dominance x additive value is

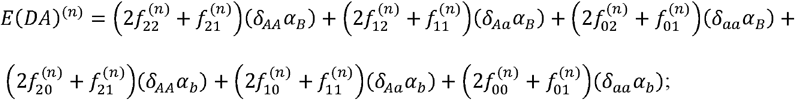

this expectation is zero only if F = 0 or if there is no LD.
6. the expectation of the dominance x dominance value is 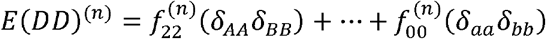; this expectation is zero only if there is no LD.

Thus, defining the additive x additive, additive x dominance, dominance x additive, and dominance x dominance epistatic values as the values expressed as deviation from its mean, *AA^(n)^ =AA - E(AA)^(n)^, AD^(n)^ = AD - E(AD)^(n)^, DA^(n)^ = DA — E(DA)^(n)^*, and *DD^(n)^ = DD — E(DD)^(n)^*, the genotypic value in an inbred population can be expressed as

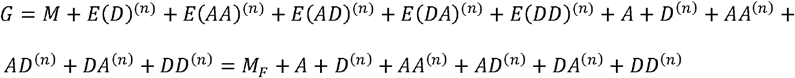

This implies that *E*(*G*) = *M_F_*. If F = 0 then

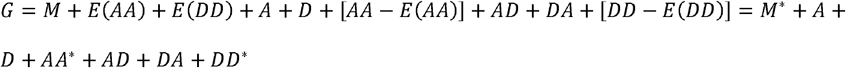

where

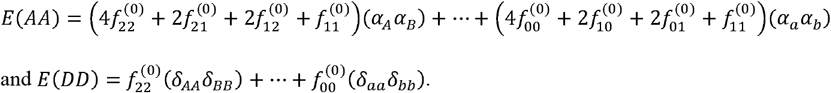

This implies that *E*(*G*) = *M_*_*. If F = 0 and there is no LD,

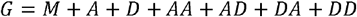

where the linear components are those defined by Kempthorne (1954). This implies that *E*(*G*) = *M*.

The assumption of LD makes very difficult to derive the components of the genotypic variance (additive, dominance, and epistatic variances and the covariances between these effects), even assuming non-inbred populations, biallelic genes, and only digenic epistasis. In respect to the types of digenic epistasis, the following can be defined (Viana 2000, 2005):

1. Complementary (*G*_22_ = *G*_21_ = *G*_12_ = *G*_11_ and *G*_20_ = *G*_10_ = *G*_02_ = *G*_01_ = *G*_00_; proportion of 9:7 in a F_2_).
2. Duplicate (*G*_22_ = *G*_21_ = *G*_20_ = *G*_12_ = *G*_11_ = *G*_10_ = *G*_02_ = *G*_01_; proportion of 15:1 in a F_2_).
3. Dominant (*G*_22_ = *G*_21_ = *G*_20_ = *G*_12_ = *G*_11_ = *G*_10_ and *G*_02_ = *G*_01_; proportion of 12:3:1 in a F_2_).
4. Recessive (*G*_22_ = *G*_21_ = *G*_12_ = *G*_11_, *G*_02_ = *G*_01_, and *G*_20_ = *G*_10_ = *G*_00_; proportion of 9:3:4 in a F_2_)
5. Dominant and recessive (*G*_22_ = *G*_21_ = *G*_12_ = *G*_11_ = *G*_20_ = *G*_10_ = *G*_00_ and *G*_02_ = *G*_01_; proportion of 13:3 in a F_2_).
6. Duplicate genes with cumulative effects (*G*_22_ = *G*_21_ = *G*_12_ = *G*_11_, and *G*_20_ = *G*_10_ = *G*_02_ = *G*_01_; proportion of 9:6:1 in a F_2_).
7. Non-epistatic genic interaction (*G*_22_ = *G*_21_ = *G*_12_ = *G*_11_, *G*_20_ = *G*_10_, and *G*_02_ = *G*_01_; proportion of 9:3:3:1 in a F_2_).

### Simulated data sets

Because the magnitude of the components of the genotypic variance generally cannot be inferred from the previous functions, all means and genetic variances and covariances were computed from the analyses of simulated data sets provided by the software *REALbreeding* (available upon request). This software uses the quantitative genetics theory that was described in the previous sections and in Viana (2004). *REALbreeding* has been used to provide simulated data in investigations in the areas of genomic selection (Viana et al. 2019), GWAS (Pereira et al. 2018), QTL mapping (Viana et al. 2017b), linkage disequilibrium (Andrade et al. 2019), population structure (Viana et al. 2013), and heterotic grouping/genetic diversity (Viana et al. 2020).

The software simulates individual genotypes for genes and molecular markers and phenotypes in three steps using user inputs. The first step (genome simulation) is the specification of the number of chromosomes, molecular markers, and genes as well as marker type and density. The second step (population simulation) is the specification of the population(s) and sample size or progeny number and size. A population is characterized by the average frequency for the genes (biallelic) and markers (first allele). The final step (trait simulation) is the specification of the individual phenotypes. In this stage, the user informs the minimum and maximum genotypic values for homozygotes, the minimum and maximum phenotypic values (to avoid outliers), the direction and degree of dominance, and the broad sense heritability. The current version allows the inclusion of digenic epistasis, gene x environment interaction, and multiple traits (up to 10), including pleiotropy. The population mean (M), additive (A), dominance (D), and epistatic (AA, AD, DA, and DD) genetic values or general and specific combining ability effects (GCA and SCA) or genotypic values (G) and epistatic values (I), depending on the population, are calculated from the parametric gene effects and frequencies and the parametric LD values. The phenotypic values (*P*) are computed assuming error effects (*E*) sampled from a normal distribution (*P = M + A + D + AA + AD+DA + DD + E = G + E* or *P = M + GCA1 + GCA2 + SCA + I + E = G + E*).

The quantitative genetics theory for epistasis does not solve the challenge of studying genetic variability and covariance between relatives in populations, using simulated data sets, even assuming simplified scenarios such as linkage equilibrium and no inbreeding. This is why there are few significant publications about the influence of epistasis on the genetic variability in populations based on the definitive theory proposed by Kempthorne (1954). Because the genotypic values for any two interacting genes are not known, there are infinite genotypic values that satisfy the specifications of each type of digenic epistasis. For example, fixing the gene frequencies (the population) and the parameters m, a, d, and d/a (degree of dominance) for each gene (the trait), the solutions *G*_22_ = *G*_21_ = *G*_12_ = *G*_11_ = 5.25 and *G*_20_ = *G*_10_ = *G*_02_ = *G*_01_ = *G*_00_ = 5.71 or *G*_22_ = *G*_21_ = *G*_12_ = *G*_11_ = 6.75 and *G*_20_ = *G*_10_ = *G*_02_ = *G*_01_ = *G*_00_ = 2.71 define complementary epistasis but the genotypic values are not the same.

The solution implemented in the software allows the user to control the magnitude of the epistatic variance (V(I)), relative to the magnitudes of the additive and dominance variances (V(A) and V(D)). As an input for the user, the software requires the ratio V(I)/(V(A) + V(D)) for each pair of interacting genes (a single value; for example, 1.0). Then, for each pair of interacting genes the software samples a random value for the epistatic value *I*_22_ (the epistatic value for the genotype AABB), assuming *I*_22_~*N*(*0,V(I)*) For complementary epistasis, for example, the other epistatic effects and genotypic values are computed as described below, checking if the genotypic values meet the specifications that were previously defined:

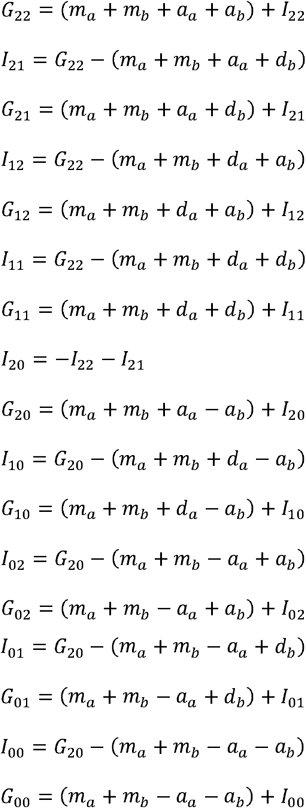

We simulated grain yield assuming 400 genes in 10 chromosomes of 200 and 50 cM (40 genes/chromosome). The average density was approximately one gene each five and one cM, respectively. We generated five populations, two with high LD level and one with low LD level, all three with an average minor allelic frequency (maf) of 0.5, and two populations with intermediate LD level and an average maf of 0.3 but with contrasting average frequency for the favorable genes (0.3/not improved and 0.7/improved). We defined positive dominance (average degree of dominance of 0.6), maximum and minimum genotypic values for homozygotes of 160 and 30 g.plt^-1^, and maximum and minimum phenotypic values of 180 and 10 g.plt^-1^. The broad sense heritability was 20%. For each population we assumed additive-dominant model and additive-dominant with digenic epistasis model, defining 100% and 30% of interacting genes and a ratio V(I)/(V(A) + V(D)) equal to 1.0. With epistasis, we assumed a single type or an admixture of the seven types. We ranged the degree of inbreeding from 0.0 to 1.0, assuming 10 generations of selfing. We also assumed 10 generations of random crosses. The population size was 5,000 per generation.

The characterization of the LD in the populations was based on the parametric D, r^2^, and D’ values for the 40 genes in chromosome 1, which were provided by *REALbreeding* (it should be similar for the other chromosomes). The heatmaps were processed using the R package pheatmap.

## Results

The analysis of the parametric LD in the populations shows that the LD level depends mainly on the gene density (Figure 1). The higher LD level was observed under high gene density (one gene each cM). Regardless of the gene density, the LD level is generally higher for the closest genes. As expected, 10 generations of random cross significantly decreased the LD level of the populations. The decrease was higher for the density of one gene each five cM, regardless of the population (on average, approximately −95% for r^2^). The average decrease in the r^2^ for the density of one gene each cM was −81%. The LD level showed only a slight decrease after 10 generations of selfing (on average, approximately −14% for r^2^, regardless of the population).

**Figure 1.**
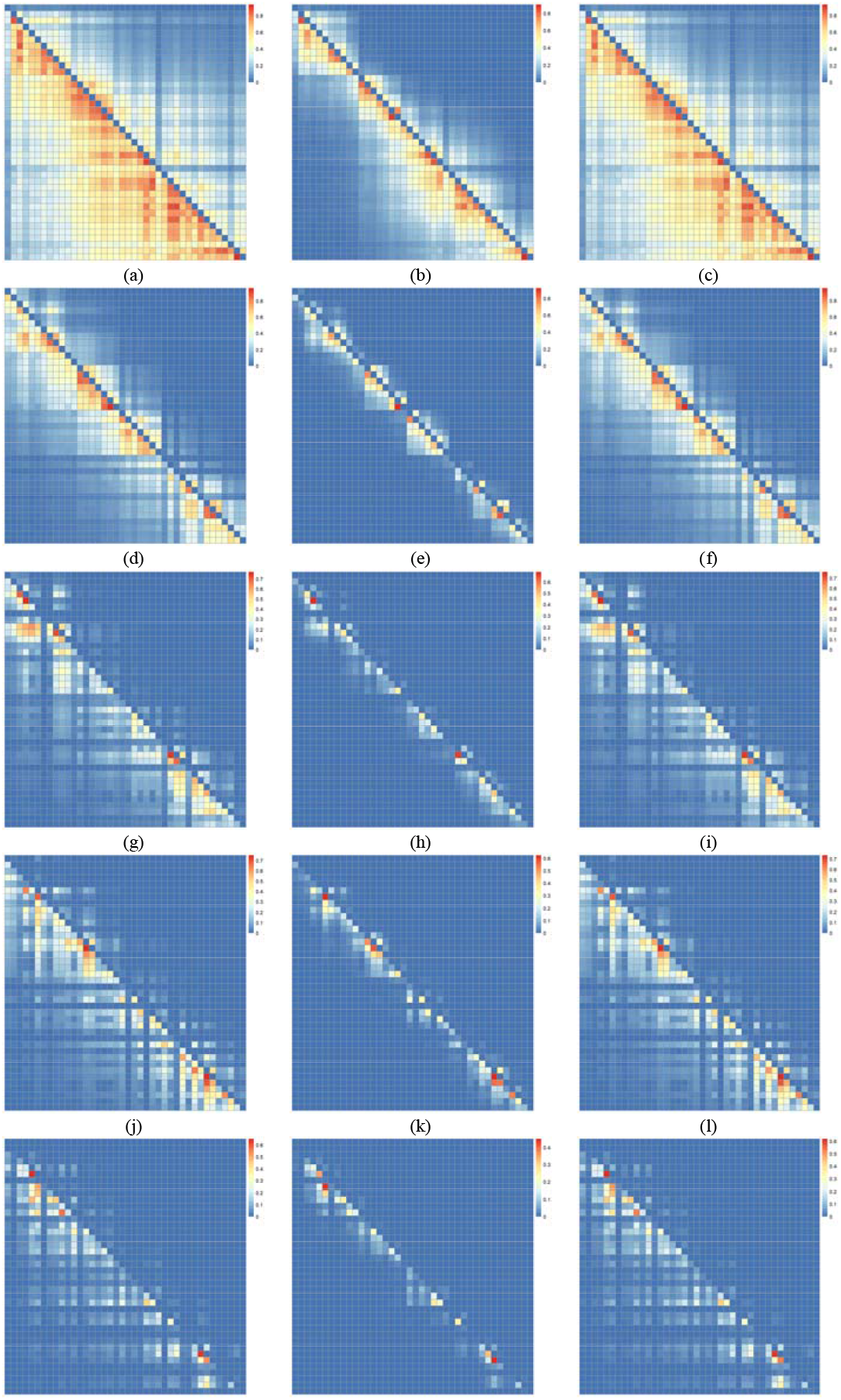
Parametric r^2^ (above the diagonal) and |D’| (below the diagonal) values for 40 genes along chromosome 1 in the populations with higher gene density and high LD (a, b, c), and lower gene density and high (d, e, and f), intermediate (g, h, i, j, k, l), and low (m, n, and o) LD levels, in the generations 0 (a, d, g, j, and m) and 10, assuming random cross (b, e, h, k, and n) or selfing (c, f, i, l, and o).

To characterize the magnitude of the genotypic variance components in non-inbred and inbred populations with contrasting LD levels, assuming no epistasis, we assumed a density of one gene each five cM. Because the populations with high and low LD levels have an average maf of 0.5, the decrease in the population mean due to inbreeding and the genotypic and additive variances are maximized, relative to populations with average maf lower or higher than 0.5. The same is true for the dominance variance in non-inbred populations. After 10 generations of selfing, the decreases in the population means were −15 and −17% for the populations with low and high LD level, respectively (Figure 2). The significance of the LD level is impressive on the magnitude of the genotypic and additive variances. Regardless of the LD level and the degree of inbreeding, the additive variance is the most important component of the genotypic variance,. The additive variance in the population with high LD is 6.8 times greater than the additive variance in the population with low LD in generation 0, 2.9 times greater after 10 generations of random cross, and 5.7 times greater after 10 generations of selfing. In the populations with intermediate LD level, the decreases in the population mean due to inbreeding are similar in magnitude. In both improved and not improved populations, there is also a decrease in the magnitude of the additive variance with random crosses and an increase with selfing. The additive variance is greater in the not improved population, regardless of the generation. In both populations the magnitude of the additive variance is intermediate to the values observed for the populations with high and low LD level.

**Figure 2.**
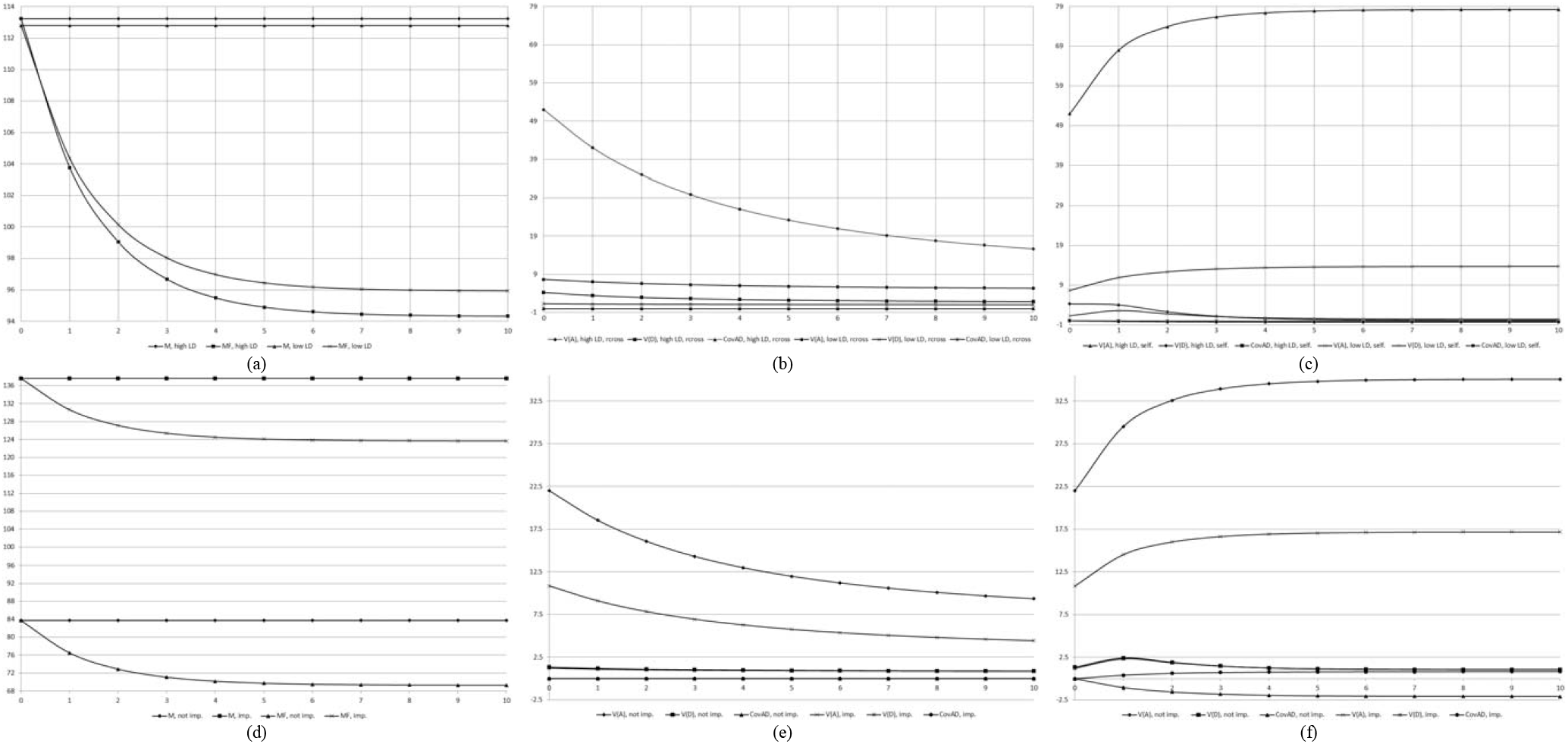
Parametric population means (M and MF; a and d), additive (V(A)) and dominance (V(D)) variances, and covariance between additive and dominance values (CovAD), in populations with high and low LD levels (b and c) and in populations with intermediate LD level (not improved and improved) (e and f), along 10 generations of random cross (b and e) and 10 generations of selfing (c and f).

To characterize the components of the genotypic variance in non-inbred and inbred populations with high LD level, assuming epistasis, we also assumed the density of one gene each five cM. Regardless of the type of epistasis and the percentage of interacting genes, there are non-significant changes in the population mean along 10 generations of random cross (remember that the average decrease in the r^2^ values was approximately −95%) (Figure 3). With 10 generations of selfing, regardless of the percentage of epistatic genes, except for recessive epistasis, the inbreeding decreased the population mean in −19 to −20% with dominant epistasis to −9 to −10% with dominant and recessive epistasis (remember that the decrease assuming no epistasis was −17%).

**Figure 3.**
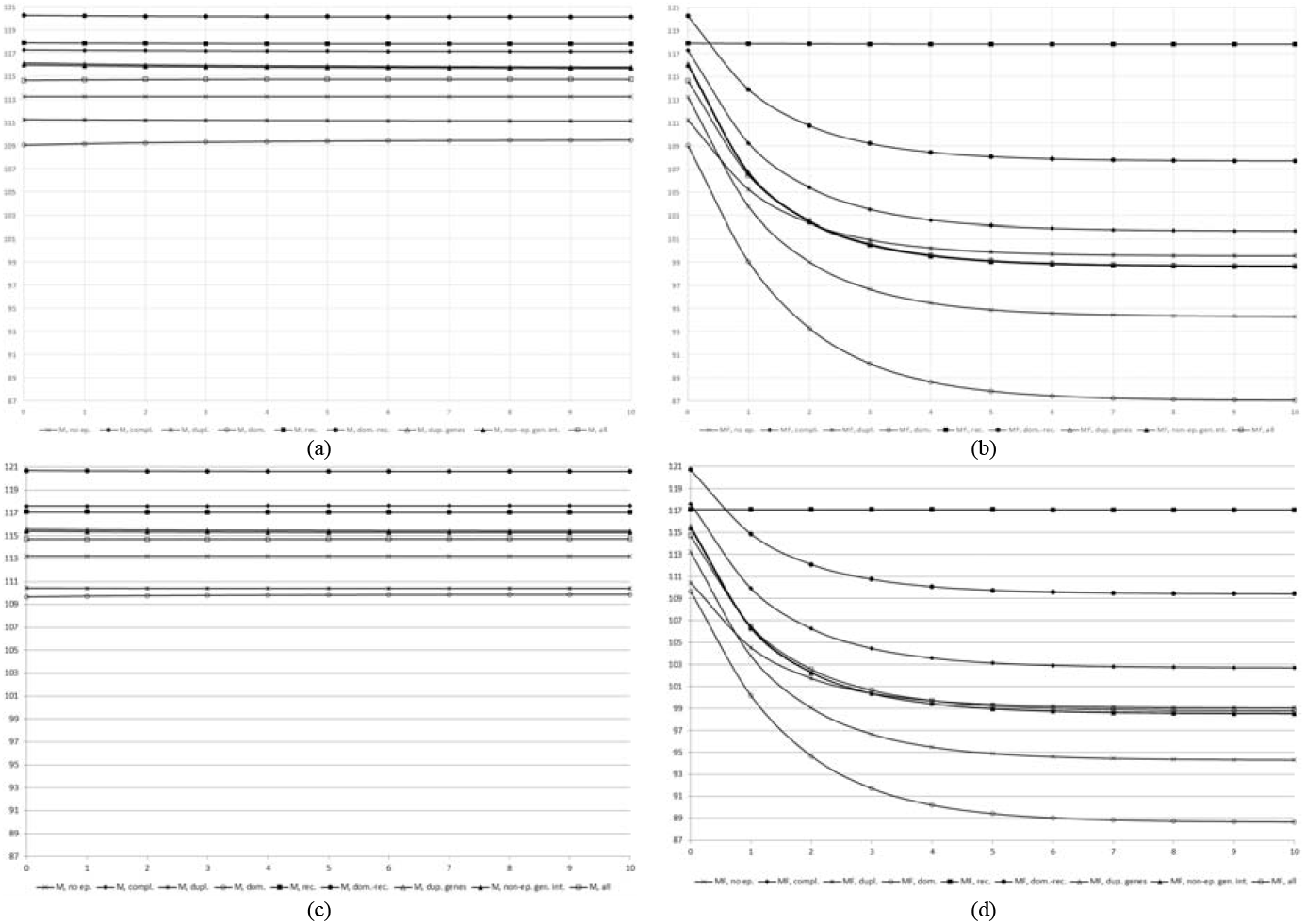
Parametric population means (M and MF) in population with high LD level, along 10 generations of random cross (a and c) and 10 generations of selfing (b and d), assuming no epistasis, a single type of digenic epistasis (complementary, duplicate, dominant, recessive, dominant and recessive, duplicate genes with cumulative effects, and non-epistatic genic interaction) or an admixture of the seven digenic types (all), and 100 (a and b) and 30% (c and d) of epistatic genes.

Regardless of the type of epistasis, the ratio epistatic variance/genotypic variance is proportional to the percentage of the epistatic genes. The epistatic variance in generation 0 corresponded to 2 to 7% (dominant epistasis) of the genotypic variance with 30% of epistatic genes, but it corresponded to 10 to 64% (duplicate epistasis) assuming 100% of epistatic genes (Figures 4 to 11). Irrespective of the type of epistasis and the percentage of epistatic genes, after 10 generations of random cross or selfing the ratio epistatic variance/genotypic variance increased in the range of 11 to 660%. This occurred because the decrease in the genotypic variance was much higher than the decrease in the epistatic variance with random cross and because the increase in the genotypic variance was much lower than the increase in the epistatic variance with selfing. With three exceptions, regardless of the type of epistasis and the percentage of epistatic genes, the most important component of the genotypic variance is also the additive variance. The magnitude of the additive variance decreased with random cross and increased with selfing. With duplicate epistasis and 100% of epistatic genes, the additive x additive variance was higher than the additive variance, throughout 10 generations of random cross or selfing. Assuming dominant epistasis and dominant and recessive epistasis, 100% of epistatic genes, and inbred population, the sum of several epistatic covariances (after four selfings) and the additive x additive variance were greater than the additive variance, respectively. Except for duplicate genes with cumulative effects and non-epistatic genic interaction, the magnitude of the additive variance was two to 16 times greater assuming 30% of epistatic genes, compared with 100% of epistatic genes, for both random cross and selfing. Assuming duplicate genes with cumulative effects and non-epistatic genic interaction, the magnitude of the additive variance was 1.1 and 1.4 times higher with 100% of interacting genes, respectively.

**Figure 4.**
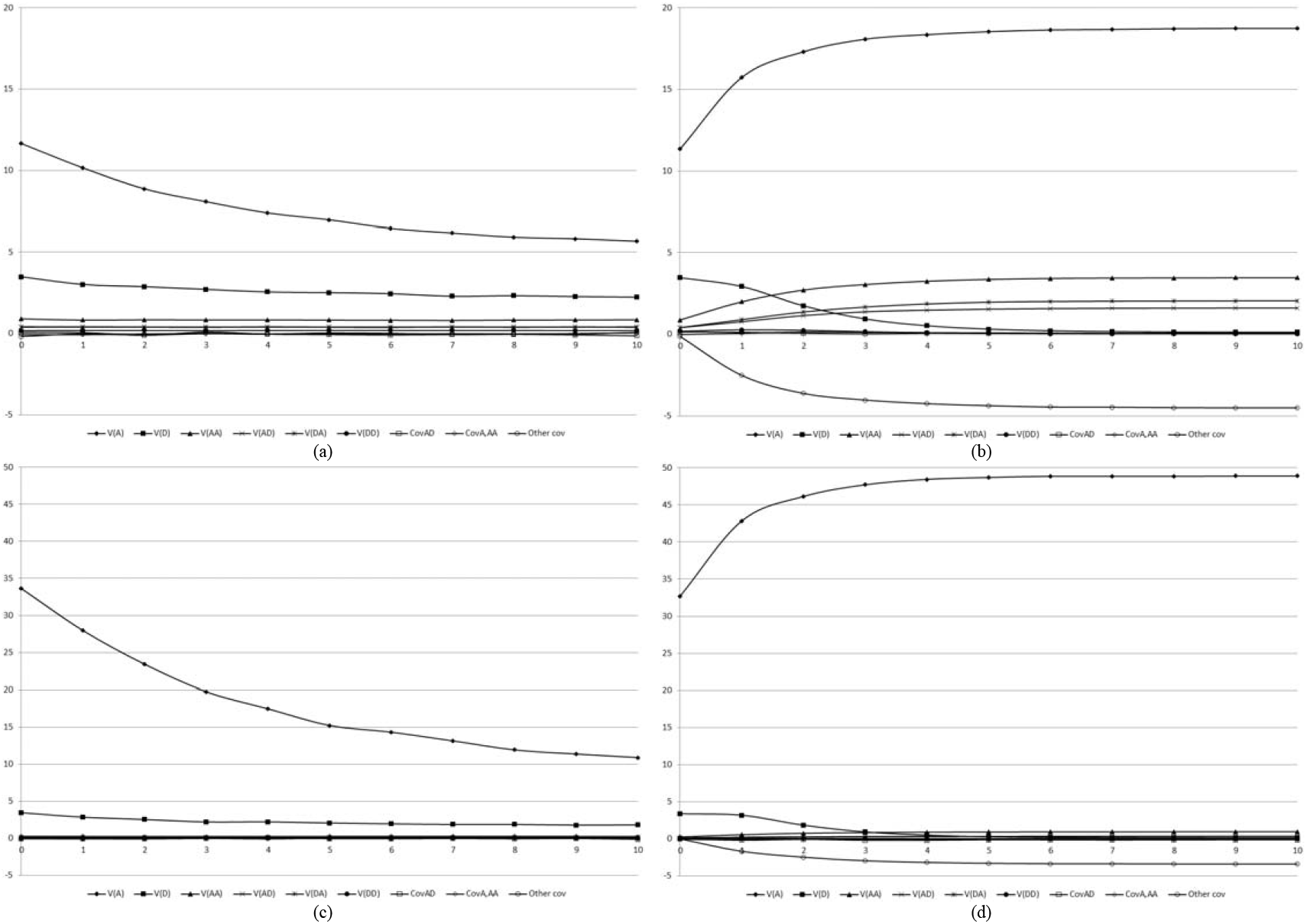
Components of the genotypic variance in population with high LD level, along 10 generations of random cross (a and c) or selfing (b and d), assuming complementary epistasis, 100 (a and b) and 30% (c and d) of epistatic genes, and sample size of 5,000 per generation.

**Figure 5.**
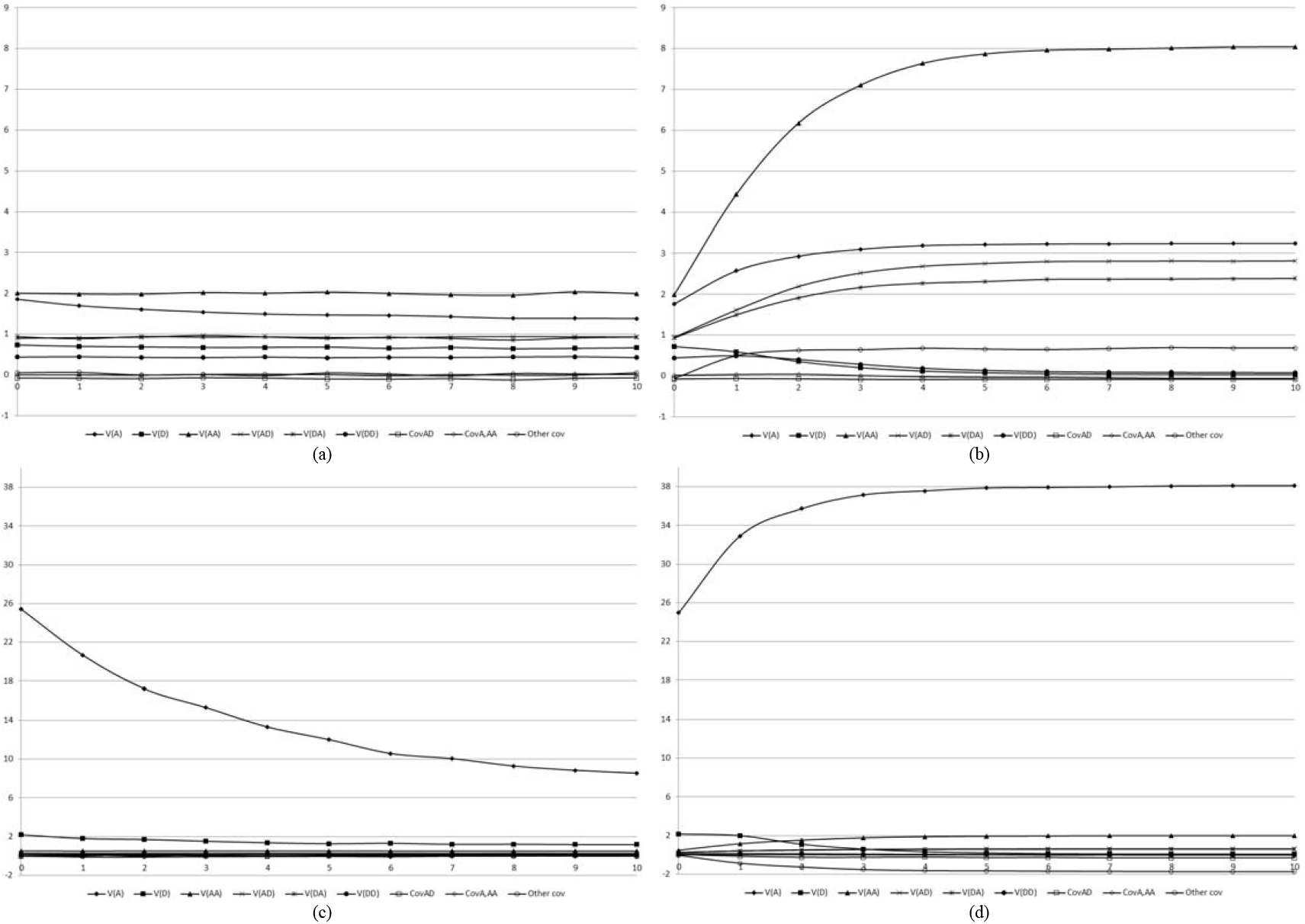
Components of the genotypic variance in population with high LD level, along 10 generations of random cross (a and c) or selfing (b and d), assuming duplicate epistasis, 100 (a and b) and 30% (c and d) of epistatic genes, and sample size of 5,000 per generation.

**Figure 6.**
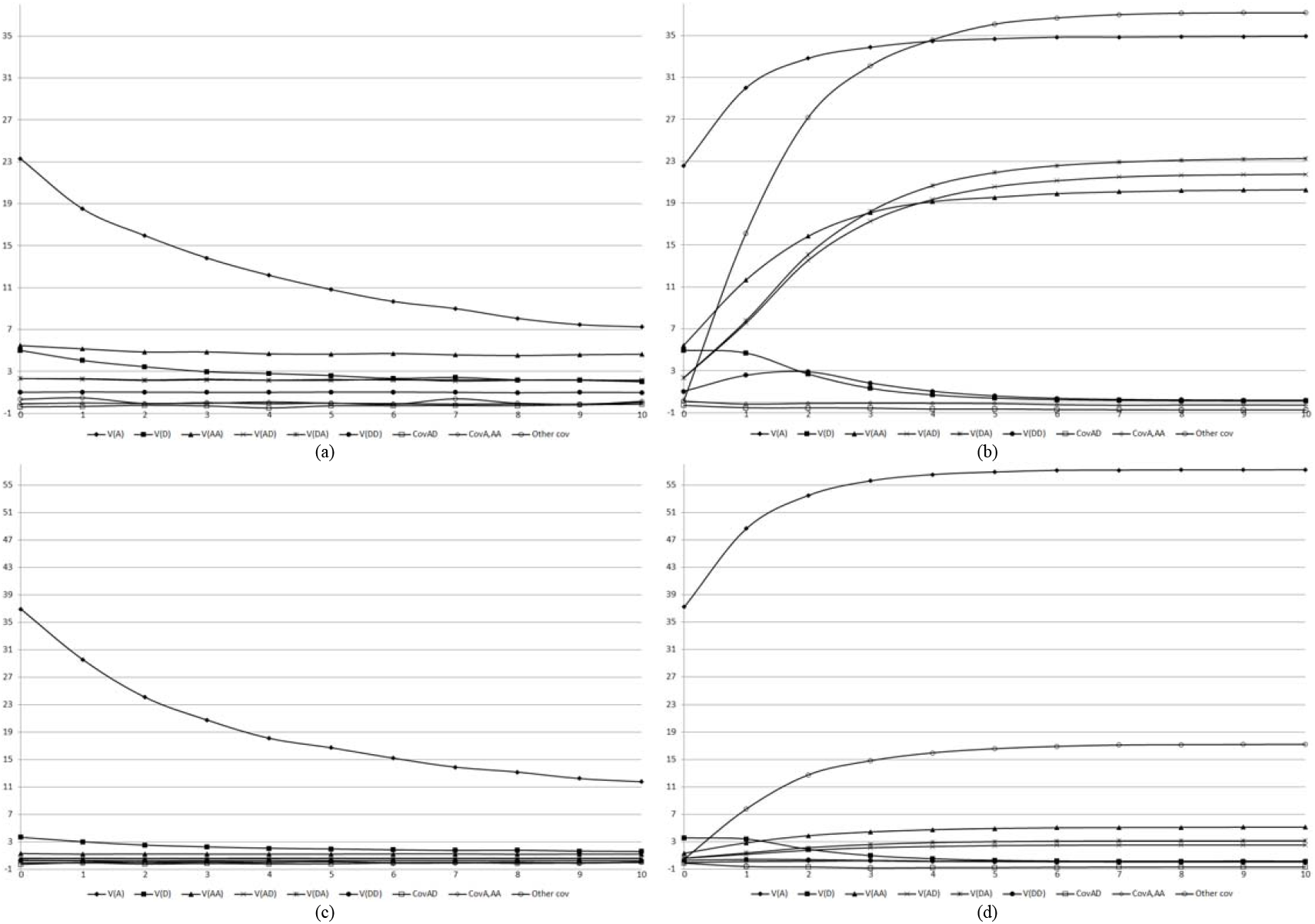
Components of the genotypic variance in population with high LD level, along 10 generations of random cross (a and c) or selfing (b and d), assuming dominant epistasis, 100 (a and b) and 30% (c and d) of epistatic genes, and sample size of 5,000 per generation.

**Figure 7.**
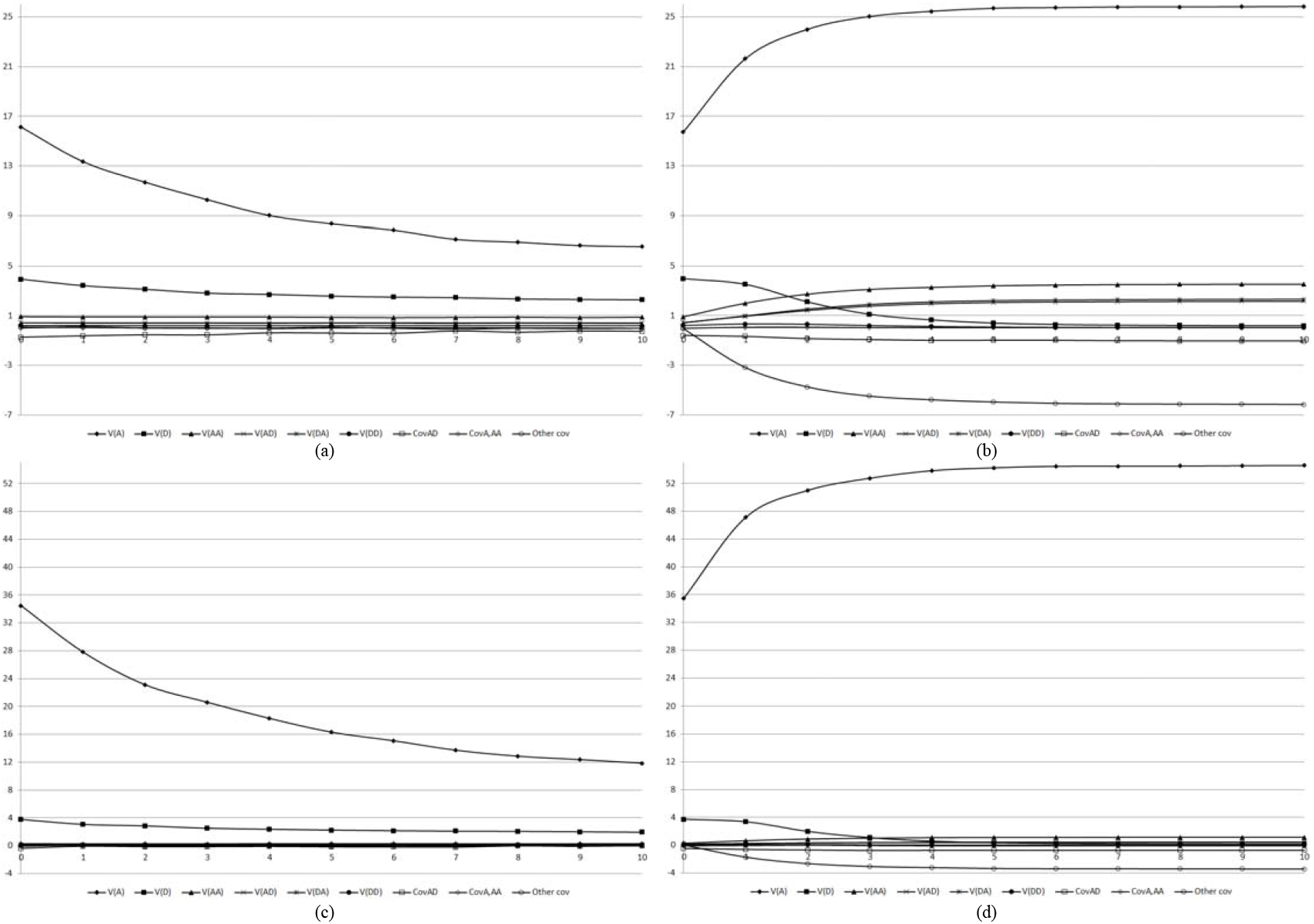
Components of the genotypic variance in population with high LD level, along 10 generations of random cross (a and c) or selfing (b and d), assuming recessive epistasis, 100 (a and b) and 30% (c and d) of epistatic genes, and sample size of 5,000 per generation.

**Figure 8.**
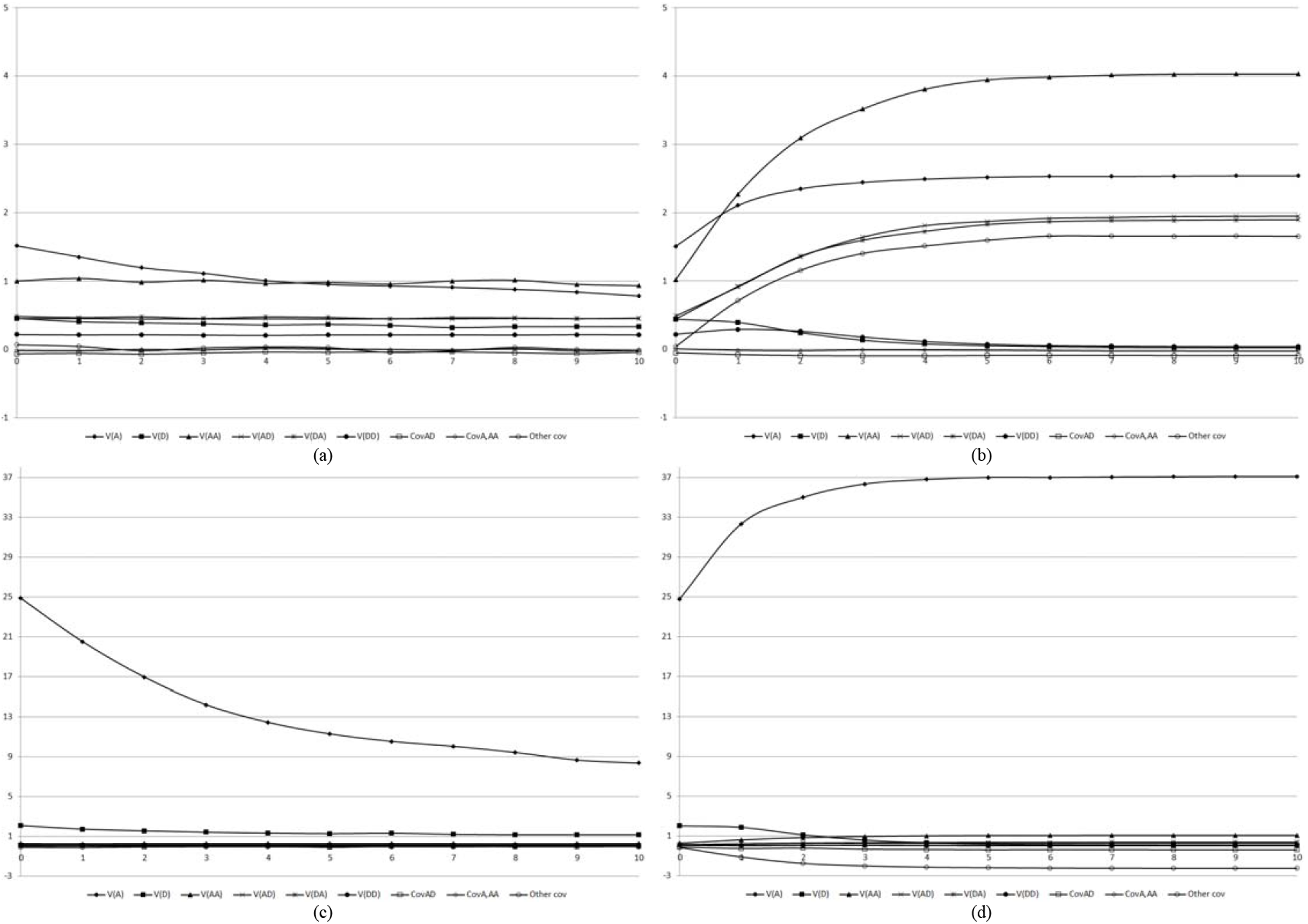
Components of the genotypic variance in population with high LD level, along 10 generations of random cross (a and c) or selfing (b and d), assuming dominant and recessive epistasis, 100 (a and b) and 30% (c and d) of epistatic genes, and sample size of 5,000 per generation.

**Figure 9.**
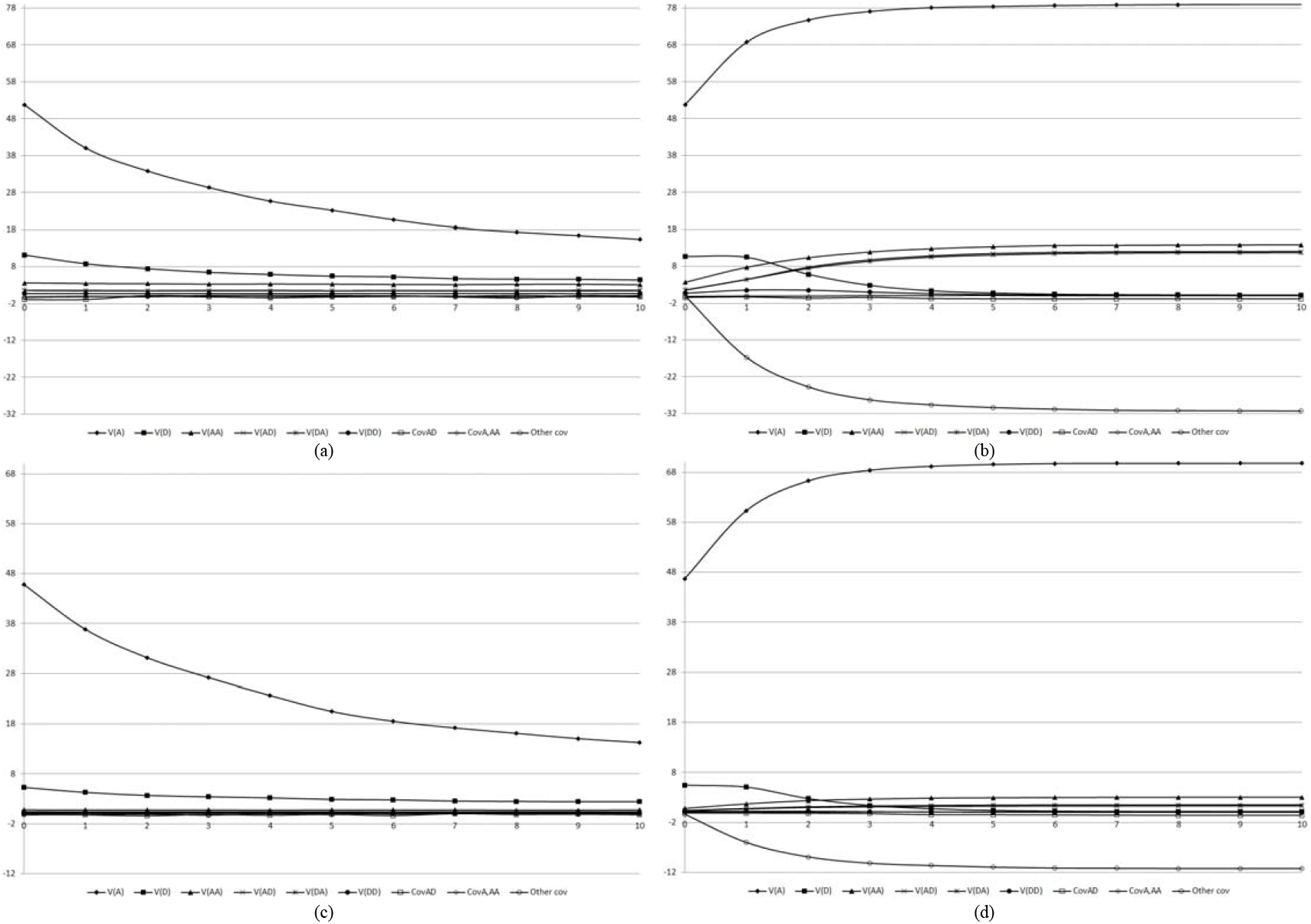
Components of the genotypic variance in population with high LD level, along 10 generations of random cross (a and c) or selfing (b and d), assuming duplicate genes with cumulative effects, 100 (a and b) and 30% (c and d) of epistatic genes, and sample size of 5,000 per generation.

**Figure 10.**
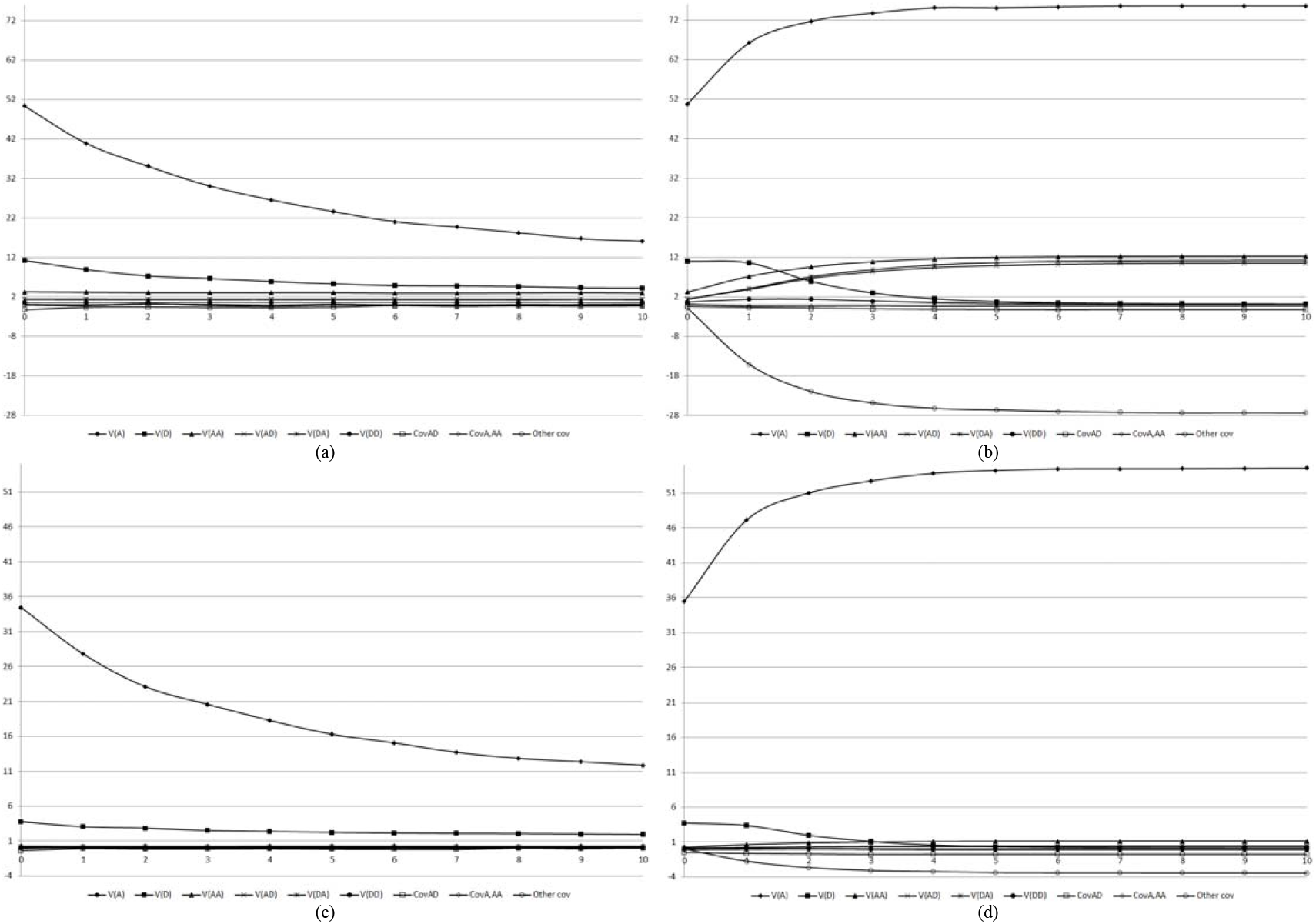
Components of the genotypic variance in population with high LD level, along 10 generations of random cross (a and c) or selfing (b and d), assuming non-epistatic genic interaction, 100 (a and b) and 30% (c and d) of epistatic genes, and sample size of 5,000 per generation.

**Figure 11.**
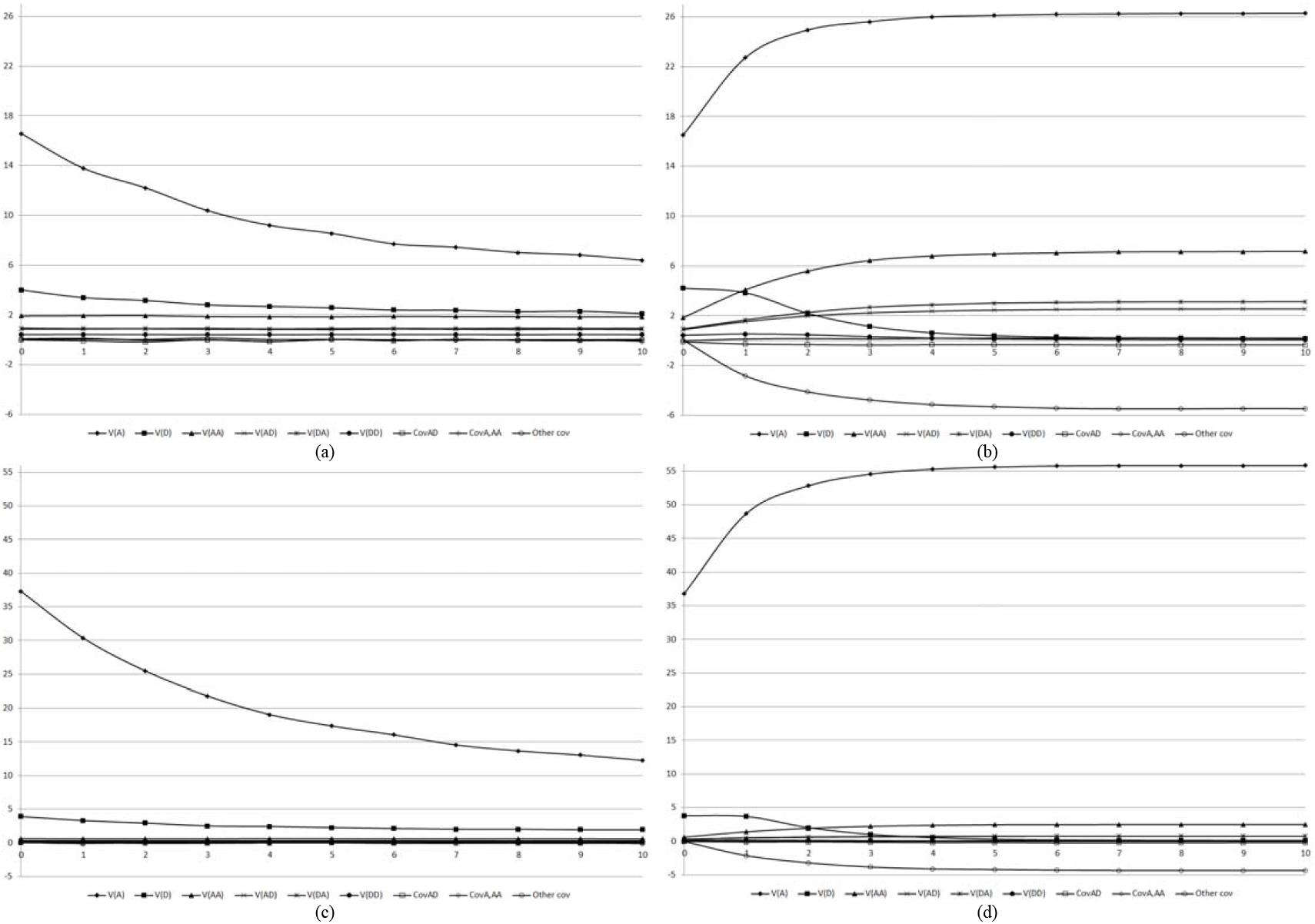
Components of the genotypic variance in population with high LD level, along 10 generations of random cross (a and c) or selfing (b and d), assuming an admixture of digenic epistasis, 100 (a and b) and 30% (c and d) of epistatic genes, and sample size of 5,000 per generation.

For the epistatic variances, except for duplicate epistasis and dominant and recessive epistasis, 100% of epistatic genes, their magnitudes are much lower than the magnitude of the additive variance. The additive x additive variance is the most important epistatic variance. Generally, a non-significant variation in the magnitudes of the epistatic variances was observed throughout 10 generations of random cross, regardless of the type of epistasis and the percentage of the epistatic genes. A significant increase in the magnitude of the additive x additive, additive x dominant, and dominant x additive variances occurred with selfing, regardless of the percentage of epistatic genes and the type of epistasis. When inbreeding increased, the dominant x dominant variance decreased in the population with high LD but increased in the other populations.

For the populations with intermediate and low LD levels, the previous inferences generally holds but the magnitudes of the genotypic and genetic variances are generally lower than the values for the population of high LD level, regardless of generation, type of epistasis, and percentage of epistatic genes, as exemplified assuming 30% of epistatic genes showing all types of epistasis (Figure 12). With no exception, the additive variance is also the most important component of the genotypic variance, regardless of the generation.

**Figure 12.**
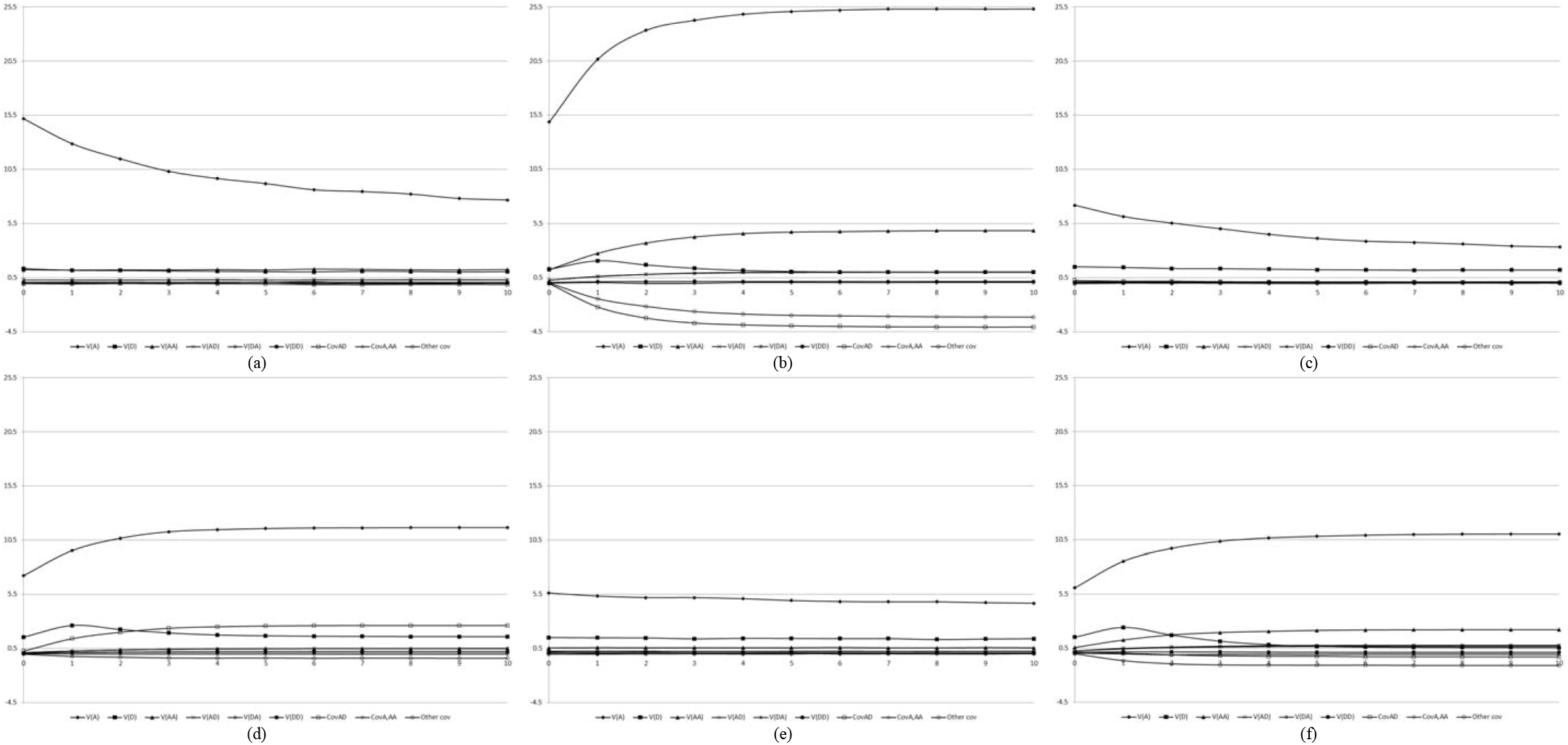
Components of the genotypic variance in the populations not improved (a and b) and improved (c and d), with intermediate LD level, and in the population with low LD level (e and f), along 10 generations of random cross (a, c, and e) or selfing (b, d, and f), assuming an admixture of digenic epistasis, 30% of epistatic genes, and sample size of 5,000 per generation.

## Discussion

Comstock and Robinson (1948) provided the basic knowledge on Quantitative Genetics, partitioning the genotypic variance in the variance due to the average effects of the genes (additive) and the variance due to allelic interaction (dominance). Assuming effects between non-allelic genes, Kempthorne (1954) and Cockerham (1954) demonstrated that the genotypic variance and the covariance between relatives depends on the additive, dominance, and four epistatic variances, assuming digenic epistasis. Cockerham and Weir (1977) derived the very complex functions for the components of the genotypic variance assuming a two-gene model with inbreeding, LD, and epistasis.

Meuwissen et al. (2001) proposed a method for predicting the additive values of non-phenotyped individuals who were genotyped for thousands of SNPs, from the model fitted by analyzing a limited number of phenotyped and genotyped individuals. Since then, the knowledge on gene effects, LD, and covariance between relatives has been directly or implicitly used for modelling additive, dominance, and epistatic SNP and QTL effects, deriving observed genomic relationship matrices, and assessing prediction accuracy, power of QTL detection, and rate of FDR in thousands of genomic prediction and association studies (Stich and Gebhardt 2011; Varona et al. 2018; Vitezica et al. 2017).

Unfortunately, for over 70 years the joint significance of LD, epistasis, and inbreeding on the genotypic variance and the covariance between relatives remained unclear. This is because the theory available, even assuming only two loci, is of “little use” (Cockerham and Weir, 1977). Additionally, no feasible theoretical model can depict the complexity of the development of a phenotype from a genotype (Robinson et al. 2014). Thus, our simulation-based study provides a better understanding of the influence of LD and epistasis on the genotypic variance and the covariance between relatives in non-inbred and inbred populations.

We assumed low to high LD levels for genes, under a relatively low gene density, and digenic epistasis. In maize, for example, the genome size is approximately 2 Gb, including approximately 42,000 to 56,000 genes. The density is approximately 1-11 genes per 100 kb over a relatively even distribution (Haberer et al. 2005). Because grain yield is affected by most of these genes, the gene density for this very complex quantitative trait should be higher than the gene density for other less complex quantitative traits. Although there is evidence for higher-order epistasis, pairwise epistasis can contribute substantially to phenotypic variation between individuals (Domingo et al. 2019).

Our main results explain the general knowledge on the efficacy of genomic prediction and GWAS provided by numerous published papers. Because the LD has a significant positive effect on the magnitude of the genotypic and genetic variances, especially the additive variance, the efficacy of the additive value genomic prediction and GWAS are proportional to the LD level for genes (Forneris et al. 2017) and to the degree of inbreeding (Martini et al. 2017). The accuracy of future predictions in non-inbred populations, such as human populations and in animal breeding, decreases mainly due to the decrease in the LD, because the covariance between relatives in distinct generations depends on the components of the genotypic variance in the first generation. Updating training data after some generations implies in defining a new training population with lower LD and, consequently, lower components of the genotypic variance, but it makes the individuals in the training and selection generations more related. However, in animal breeding, probable because changes in the SNP and QTL frequencies due to selection, increasing the LD level for SNPs and QTLs, field and simulation-based studies have showed increase in the prediction accuracy (Neyhart et al. 2017).

Because the epistatic variances have a lower magnitude relative to the additive variance, some field and simulation-based investigations have not shown an increase in the additive value prediction accuracy by fitting SNP epistatic effects (Chen et al. 2019; Forneris et al. 2017; Vitezica et al. 2018). However, including epistasis can improve accuracy and unbiasedness of genomic predictions (Su et al. 2012) as well as QTL detection power and control of the type I error in GWAS (Monir and Zhu 2017; Pecanka et al. 2017; Stich and Gebhardt 2011). However, epistatic variances can be the most important components of the genotypic variance with higher-order epistasis, especially assuming a high number of interacting genes (Viana 2000). Because the additive x additive variance is the most important epistatic variance, predicting the additive value of inbred plants over generations of selfing, including epistasis, can increase the selection efficiency (Jiang and Reif 2015).

In conclusion, our main results from a simulation-based study supported by quantitative genetics theory involving LD, epistasis, and inbreeding were: 1) the LD level for genes, even under a relatively low gene density, has a significant positive effect on the magnitude of the components of the genotypic variance in non-inbred and inbred populations; 2) assuming digenic epistasis, the additive variance is the most important component of the genotypic variance in non-inbred and inbred populations; 3) the magnitude of the epistatic variance is proportional to the percentage of interacting genes, regardless of the degree of inbreeding; 4) except for duplicate and dominant and recessive epistasis, with 100% of epistatic genes, the additive variance is greater than the epistatic variance; and 5) regardless of the degree of inbreeding, the additive x additive variance is the most important component of the epistatic variance. This explains why LD for genes and relationship information are key factors affecting the genomic prediction accuracy of complex traits and the efficacy of GWAS.

## Acknowledgements

We thank the National Council for Scientific and Technological Development (CNPq), the Brazilian Federal Agency for Support and Evaluation of Graduate Education (Capes; Finance Code 001), and the Foundation for Research Support of Minas Gerais State (Fapemig) for financial support.

## Data Availability

The dataset is available at https://doi.org/10.6084/m9.figshare.13607306.v1.

## Conflict of Interest

The authors declare that they have no conflicts of interest.

## Author Contributions

Both authors contributed equally.

## Notes

https://doi.org/10.6084/m9.figshare.13607306.v1

### Competing Interest Statement

The authors have declared no competing interest.

